# CGAT-core: a python framework for building scalable, reproducible computational biology workflows

**DOI:** 10.1101/581009

**Authors:** Adam P Cribbs, Sebastian Luna-Valero, Charlotte George, Ian M Sudbery, Antonio J Berlanga-Taylor, Steven N Sansom, Thomas Smith, Nicholas E. Ilott, Jethro Johnson, Jakub Scaber, Katherine Brown, David Sims, Andreas Heger

## Abstract

In the genomics era computational biologists regularly need to process, analyse and integrate large and complex biomedical datasets. Analysis inevitably involves multiple dependent steps, resulting in complex pipelines or workflows, often with several branches. Large data volumes mean that processing needs to be quick and efficient and scientific rigour requires that analysis be consistent and fully reproducible. We have developed CGAT-core, a python package for the rapid construction of complex computational workflows. CGAT-core seamlessly handles parallelisation across high performance computing clusters, integration of Conda environments, full parameterisation, database integration and logging. To illustrate our workflow framework, we present a pipeline for the analysis of RNAseq data using pseudoalignment.

**Availability:** CGAT-core is freely available under an MIT licence for installation and use, including source code at https://github.com/cgat-developers/cgat-core

**Contact:** (AH), (DS), (AC)

***Supplementary information***:

## 1. Introduction

Genomic technologies have given researchers the ability to produce large amounts of data at relatively low cost. Bioinformatic analyses typically involve passing data through a series of manipulations and transformations, called a pipeline or workflow. The need for tools to manage workflows is well established, with a wide range of options available from graphical user interfaces such as Galaxy (Afgan, et al., 2016) and Taverna (Wolstencroft, et al., 2013), aimed at non-programmers, to Snakemake, Nextflow, Toil, Ruffus and others (Di Tommaso, et al., 2017; Fisch, et al., 2015; Gafni, et al., 2014; Golosova, et al., 2014; Koster and Rahmann, 2012; Nocq, et al., 2013; Okonechnikov, et al., 2012; Vivian, et al., 2017) developed with computational biologists in mind. These tools differ in their portability, scalability, parameter handling, extensibility, and ease of use. In a recent survey (Leipzig, 2017), the tool rated highest for ease of pipeline development was Ruffus (Goodstadt, 2010), a Python package that wraps pipeline steps in discrete Python functions, called ‘tasks’. It uses Python decorators to track the dependencies between tasks, ensuring that dependent tasks are completed in the correct order and independent tasks can be run in parallel. If a pipeline is interrupted before completion, or new input files are added, only data sets that are missing or out-of-date are rerun. Ruffus implements a wide range of decorators that allow complex operations on input files including: conversion of a single input file to a single output file; splitting of a file into multiple files (and vice versa) and conditional merging of multiple input files into a smaller number of outputs. More advanced options include combining combinations or permutations of input files and conditional execution based on input parameters. Use of decorators means that Ruffus pipelines are native Python scripts, rather than the domain specific languages (DSLs) used in other workflow tools. A key advantage of this, in addition to python being an already widely understood language in computational biology, is that individual steps can use arbitrary python code, both in how they are linked together and in the actual processing task.

Here, we introduce Computational Genomics Analysis Toolkit (CGAT)-core, an open-source python library that extends the functionality of Ruffus by adding cluster interaction, parameterisation, logging, database interaction and Conda (https://conda.io) environment switching.

## 2. Main features

CGAT-core extends the functionality of Ruffus by providing a common interface to control distributed resource management systems using DRMAA (Distributed Resource Management Application API, http://www.drmaa.org/). Currently, we support interaction with Sun Grid Engine, Slurm and PBS-pro/Torque. The execution engine enables tasks to be run locally or on a high-performance computing cluster and supports cluster distribution of both command line scripts (*cgatcore.run*) and python functions (*cgatcore.cluster*). System resources (number of cores to use, amount of RAM to allocate) can be set on a per-pipeline, per-task, or per task-instance basis, even allowing allocation to be based in variables, for example input file size.

The parameter management component encourages the separation of workflow/tool configuration from implementation to build re-usable workflows. Algorithm parameters are collected in a single human-readable yaml configuration file. Thus, parameters can be set specifically for each dataset, without the need to modify the code. For example, sequencing data can be aligned to a different reference genome, by simply changing the path to the genome index in the yaml file. Both pipeline-wide and job-local parameters are automagically substituted into command line statements at execution-time.

To assist with reproducibility, record keeping and error handling CGAT-core provides multilevel logging during workflow execution, recording full details of runtime parameters, environment configuration and tracking job submissions. Additionally, CGAT-core provides a simple, lightweight interface for interacting with relational databases such as SQLite (*cgatcore.database*), facilitating loading of analysis results at any step of the workflow, including combining output from parallel steps in single wide- or long-format tables.

CGAT-core can load a different Conda environment for each step of the analysis, enabling the use of tools with conflicting software requirements. Furthermore, providing Conda environment files alongside pipeline scripts ensures that analyses can be fully reproduced.

CGAT-core workflows are Python scripts, and as such are stand-alone command line utilities that do not require the installation of a dedicated service. In order to reproducibly execute our workflows, we provide utility functions for argument parsing, logging and record keeping within scripts (*cgatcore.experiment*). Workflows are started, inspected and configured through the command line interface. Therefore, workflows become just another tool and can be re-used within other workflows. Furthermore, workflows can leverage the full power of Python, making them completely extensible and flexible.

## 3. Implementation

To illustrate a simple use case of CGAT-core, we have built an example RNAseq analysis pipeline, which performs read counting using Kallisto (Bray, et al., 2016) and differential expression using DESeq2 (Love, et al., 2014). This workflow and Conda environment are contained within our CGAT-showcase repository (https://github.com/cgat-developers/cgat-showcase). The workflow highlights how simple pipelines can be constructed using CGAT-core, demonstrating how the pipeline can be configured using a yaml file, how third-party tools can be executed efficiently across a cluster or on a local machine, and how data can be easily loaded into a database. Furthermore, we and others have been extensively using CGAT-core to build pipelines for computational genomics (https://github.com/cgat-developers/cgat-flow).

CGAT-core is implemented in Python 3 and installable via Conda and PyPI (https://pypi.org) with minimal dependencies. We have successfully deployed and tested the code on OSX, Red Hat and Ubuntu. We have made CGAT-core and associated repositories open-source under the MIT licence, allowing full and free use for both commercial and non-commercial purposes. Our software is fully documented (http://cgat-core.readthedocs.io/), version controlled and has extensive testing using continuous integration (https://travis-ci.org/cgat-developers.) We welcome community participation in code development and issue reporting through GitHub.

## 4. Discussion

CGAT-core extends the popular Python workflow engine Ruffus by adding desirable features from a variety other workflow systems to form an extremely simple, flexible and scalable package. CGAT-core provides seamless high-performance computing cluster interaction and adds Conda environment integration for the first time. In addition, our framework focuses on simplifying the pipeline development and testing process by providing convenience functions for parameterisation, database interaction, logging and pipeline interaction.

The ease of pipeline development enables CGAT-core to bridge the gap between exploratory data analysis and building production workflows. A guiding principle is that it should be as easy (or easier) to complete a series of tasks using a simple pipeline compared to using an interactive prompt, especially once cluster submission is considered. CGAT-core enables the production of analysis pipelines that can easily be run in multiple environments to facilitate sharing of code as part of the publication process. Thus, CGAT-core encourages a best-practice reproducible research approach by making it the path of least resistance. For example, exploratory analysis in Jupyter Notebooks can be converted to a Python script or used directly in the pipeline. Similarly, exploratory data analysis in R, or any other language, can easily be converted to a script that can be run by the pipeline. This lightweight wrapping of quickly prototyped analysis forms a lab book, enabling rapid reproduction of analyses and reuse of code for different data sets.

## Availability and requirements

**Project Name**: CGAT-core

**Project Home Page**: http://cgat-core.readthedocs.io/

**Operating system(s)**: Linux/Unix

**Programming language**: Python 3

**Other requirements**:

**Licence**: MIT

**Any restrictions to use by non-academics**: None

## Abbreviations

DSL: Domain Specific Languages
API: Application Programming Interface
DRMAA: Distributed Resource Management Application API

## Competing interests

The authors declare that they have no competing interests.

